# Integrated sequence and gene expression analysis of mouse models of breast cancer reveals critical events with human parallels

**DOI:** 10.1101/375154

**Authors:** Jonathan P Rennhack, Matthew Swiatnicki, Yueqi Zhang, Caralynn Li, Evan Bylett, Christina Ross, Karol Szczepanek, William Hanrahan, Muthu Jayatissa, Kent Hunter, Eran R. Andrechek

**Author notes:** Corresponding author, Eran R. Andrechek, Department of Physiology Michigan State University, 2194 BPS Building, 567 Wilson Road, East Lansing, MI, 48824, (517)884-5042 Office, (517)884-5020 Lab, (517)355-5125.

## Abstract

Mouse models have an essential role in cancer research, yet little is known about how various models resemble human cancer at a genomic level. However, the shared genomic alterations in each model and corresponding human cancer are critical for translating findings in mice to the clinic. We have completed whole genome sequencing and transcriptome profiling of two widely used mouse models of breast cancer, MMTV-Neu and MMTV-PyMT. This genomic information was integrated with phenotypic data and CRISPR/Cas9 studies to understand the impact of key events on tumor biology. Despite the engineered initiating transgenic event in these mouse models, they contain similar copy number alterations, single nucleotide variants, and translocation events as human breast cancer. Through integrative in vitro and in vivo studies, we identified copy number alterations in key extracellular matrix proteins including Collagen 1 Type 1 alpha 1 (Col1a1) and Chondroadherin (CHAD) that drive metastasis in these mouse models. Importantly this amplification is also found in 25% of HER2+ human breast cancer and is associated with increased metastasis. In addition to copy number alterations, we observed a propensity of the tumors to modulate tyrosine kinase mediated signaling through mutation of phosphatases. Specifically, we found that 81% of MMTV-PyMT tumors have a mutation in the EGFR regulatory phosphatase, PTPRH. Mutation in PTPRH led to increased phospho-EGFR levels and decreased latency. Moreover, PTPRH mutations increased response to EGFR kinase inhibitors. Analogous PTPRH mutations are present in lung cancer patients and together this data suggests that a previously unidentified population of human lung cancer patients may respond to EGFR targeted therapy. These findings underscore the importance of understanding the complete genomic landscape of a mouse model and illustrate the utility this has in understanding human cancers.

## Introductory paragraph

Heterogeneity in human breast cancer is present in genomic events, gene expression, metastatic potential, and treatment response. To assess gene function in tumor biology, studies have used model systems, including genetically engineered mouse models (GEMMs). Recent work characterized the transcriptional landscape of breast cancer GEMMs, with particular attention paid to relationships with human breast cancer^1–3^. However, whether additional genomic events are required for tumor development and progression in these models remains unknown. Here we present whole genome sequencing data of two highly utilized mouse models of breast cancer, MMTV-Neu^4^ and MMTV-PyMT^5^. In PyMT tumors we identified a highly conserved mutation in the protein tyrosine phosphatase receptor (*Ptprh*) resulting in elevated EGFR activity and erlotinib sensitivity. In Neu tumors, a copy number alteration including Collagen Type 1 Alpha 1 (*Col1a1*) and Chondroadherin (*Chad*) altered metastatic potential, which was validated through genetic ablation. Together, this data demonstrates that genomic alterations beyond the initiating oncogene need to be considered when choosing a model system for breast cancer.

## Main text

To characterize the genomic landscape of the MMTV-Neu and MMTV-PyMT tumors, we created a tumor database with complete phenotypic characterization including tumor latency, histology, and metastatic burden (Table S1). Representative tumors from this database were selected for whole genome sequencing and whole transcriptome profiling by microarray. The analysis pipeline then correlated phenotypic changes with molecular profiling, including transcriptomics and sequence alterations. The resulting genes were then filtered through human breast cancer datasets to ensure relevance to human breast cancer and confirmed with *in vitro / in vivo* experiments (Figure 1A). A high degree of transcriptomic diversity both between and within each model was observed in hierarchical clustering (Figure 1B). As expected, this heterogeneity correlated with tumor histological subtype rather than tumor model, consistent with recent studies^1, 6^. It was hypothesized that these differences in expression were driven by genomic changes.

**Figure 1:**
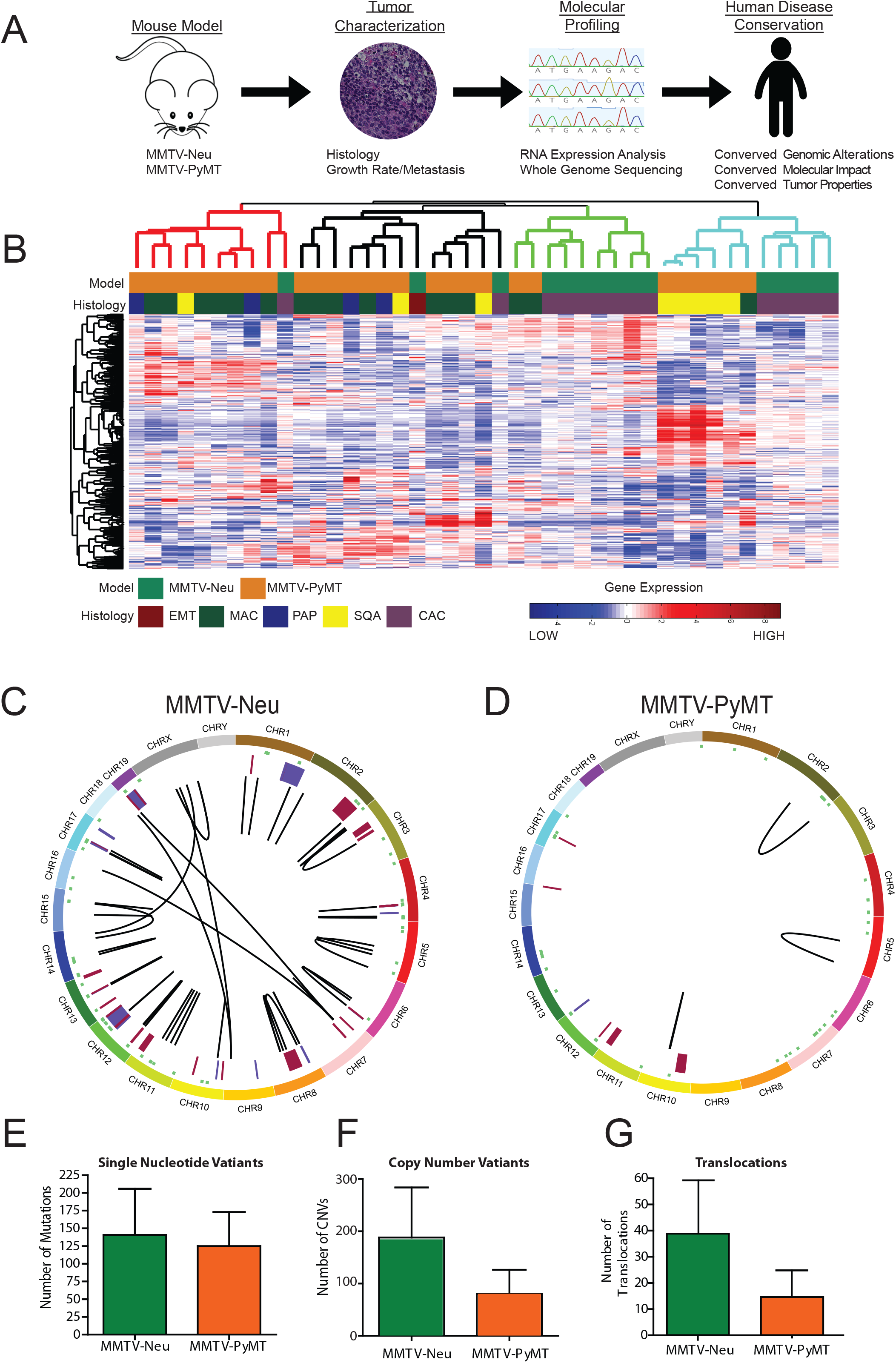
Genomic landscape of MMTV-Neu and MMTV-PyMT tumors. The schematic representation of the project workflow is depicted (A), where mammary tumors from two major mouse models are completely characterized through histological, molecular, sequence and transcriptomic methods. After data integration and analysis, the tumors were compared to human cancers at both genomic and phenotypic levels. Gene expression patterns from MMTV-Neu and MMTV-PyMT tumors were compared by unsupervised clustering, revealing substantial heterogeneity both between and within models. Tumors clustered largely based on histological subtype and not simply genotype. SQU – squamous, MAC – microacinar, PAP – papillary, and CAC – comedo-adenocarcinoma (n=15 for MMTV-Neu, n=25 for MMTV-PyMT) (B). Circos plots from whole genome sequencing results for MMTV-Neu (C) and MMTV-PyMT (D) tumors revealed differences between the strains for genomic alterations. Plots display from outside in; Chromosomal location (Each chromosome is unique color), SNVs (green), copy number alterations (Amplification – Red and Deletions – Blue), and translocations (black lines). Variation from multiple tumors is shown for Single Nucleotide Variants (E), Copy Number Variants (F) and translocations (G) (n=3 for each).

Following standard informatic pipelines, the whole genome sequence was analyzed. To validate bioinformatic calls of SNVs and CNVs we used PCR and qPCR, observing a validation rate of 85% (Table S2). Whole genome sequencing revealed large differences in the genomic landscape of the MMTV-Neu (Figure 1C) and MMTV-PyMT (Figure 1D) tumors. The two tumor models had similar numbers of SNVs (Figure 1E, Table S3), however both models were ~20X more stable than human breast tumors with 0.049 mutations/megabase in the mouse models in comparison to an average of approximately 1 mutation/megabase in breast cancer^7^. Copy number alterations (Figure 1F, Table S4) and translocations (Figure 1G, Table S5) were more frequent in the MMTV-Neu model relative to MMTV-PyMT.

To understand the specific role of copy number alterations within the two models we compared copy number variants present in the mouse models with those also in the human breast cancer. This analysis identified 11 candidate genes which were highly altered in breast cancer (Figure S1) and predicted to impact tumor biology based upon a literature screen. qPCR gene copy number analysis across an extended tumor panel (15 MMTV-PyMT, 10 MMTV-Neu) identified the rate at which each copy number variant occurred throughout the model (Figure 2A). This analysis showed that while each of the copy number variants predicted through bioinformatic means were valid (Table 2S), the depth of the amplification was largely around 1.5 fold indicating shallow amplification events (Figure 2B). Interestingly we identified the largest diversity of copy number profiles in the 11D locus. This locus includes a total of 40 genes, 19 with transcriptomic differences. Depending on the presence or absence of the locus, the tumors exhibited striking differences in structure and behavior. We identified dramatic differences in the tumors with the presence of an 11D amplification with regards to collagen content through a Mason’s trichrome (Figure 2C) stain and the presence of metastatic lesions in the lungs (Figure 2D).

**Figure 2:**
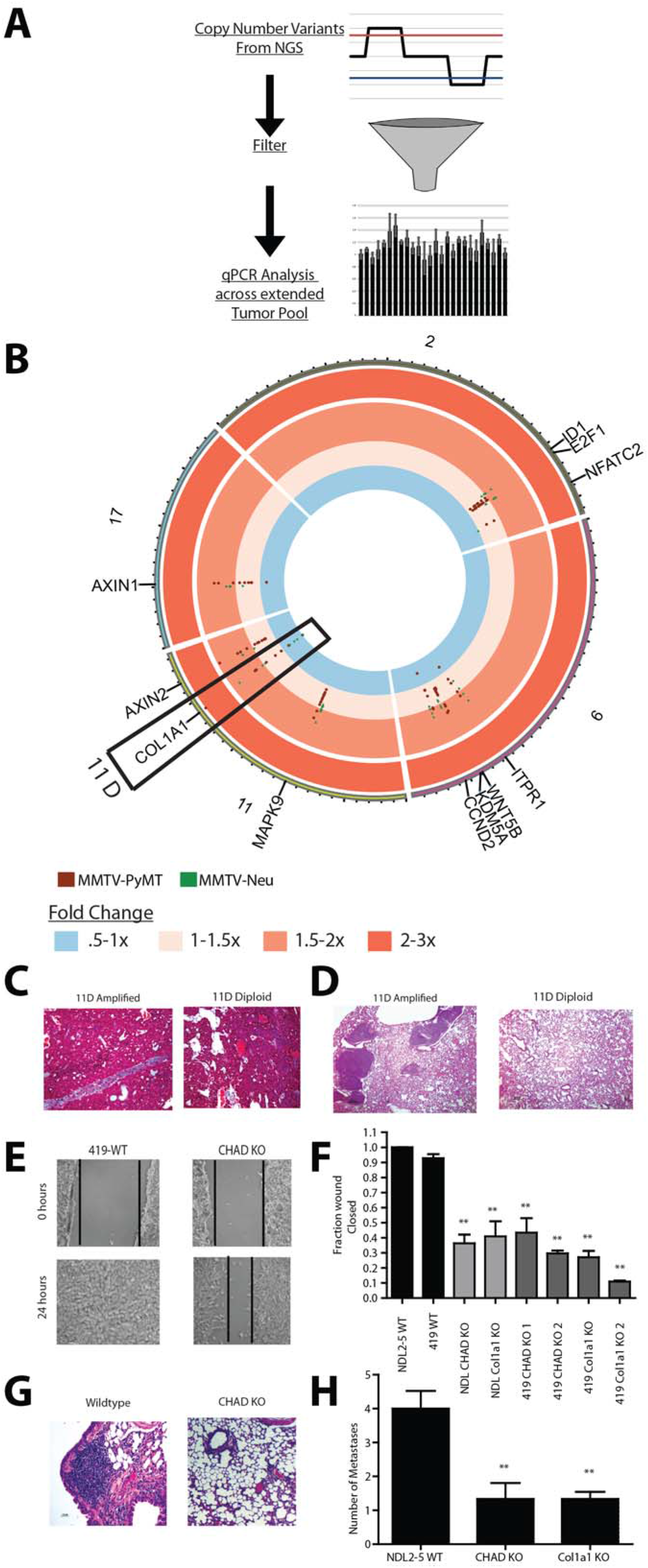
Copy number alterations alter metastatic potential. Schematic representation of filtering of copy number variants (A) where copy number variants were detected from NGS and filtered to the top 11 variants by selecting those genes that encompassed both mouse models and were conserved in human breast cancer. These genes were then assayed using qPCR analysis across a larger panel of MMTV-Neu (n=10) and MMTV-PyMT (n=15) tumors and depicted using a circos plot for genes in chromosomes 2, 6, 11 and 17 (B). Deletion on the interior of the plot to a 3 fold amplification on the exterior of the plot is shown. A key copy number alteration in the 11D region encompassing the *Col1a1* gene (boxed) was observed to correlate with reduction and lack of collagen alignment in Masons trichrome staining (C) and an increase in metastases in the lungs of mice with *Col1a1* amplification in the primary tumors (D). CRISPR-Cas9 mediated knockout of two key genes within this region, *Col1a1* and *Chad*, show defects in wound healing (E, F) (n=9). Knockout also impaired the ability to colonize the lung through a tail vein injection (G, H) (WT n=12, Chad KO n=9, Col1a1 KO n=6). (**=P<.01 for F and H)

To identify the driving genes of the metastatic phenotype associated with 11D amplification, we examined human breast cancer for distant metastasis free survival outcomes and then created CRISPR-Cas9 generated knockouts of two potential metastasis related proteins within the region, Collagen type 1 alpha 1 (*Col1a1*) and Chondroadherin (*Chad*). Knockouts were generated in two mouse driven tumor cell lines NDL2-5^8^ and PyMT 419^9^ (Figure S2). NDL2-5 is an 11D amplified Neu driven line, while the 419 line is diploid for the 11D locus and is driven by PyMT expression. Knockouts of each gene in both cell lines revealed defects in the ability to migrate in a wound healing assay (Figure 2E and Figure 2F). Migration was partially rescued with addback of wildtype *Col1a1* or *Chad*, demonstrating that migration defects were not due to off target effects (Figure S3). Defects in lung colonization in a tail vein injection were also observed with the *Col1a1* and *Chad* knockout cell lines (Figure 2G and Figure 2H).

Mouse chromosome 11D is conserved in humans and is analogous to chromosomal region 17q21.33. There is similar amplification event at 17q21.33, including *COL1A1* and *CHAD* that occurs in 8% of breast cancer patients. Array CGH from the TCGA data^10^ demonstrates that *COL1A1/CHAD* amplification was distinct from HER2 amplification (Figure 3A). Importantly, this amplification is subtype specific; 25% of Her2+ breast cancers have a co-amplification of the 17q21.33 region along with the HER2 amplicon while only 6% of Luminal A, 7% of Luminal B, and 1.2% of Basal breast cancers have amplification (Figure 3B). To investigate the transcriptional impact of the amplification event we used weighted gene correlation network analysis^11^. This identified a robust transcriptional signature that differentiated *COL1A1/CHAD*, Her2 positive tumors from Her2 positive tumors without the amplification event (Table S6). Unsupervised hierarchical clustering readily identified separation of the two HER2 positive subtypes based on this signature (Figure 3C). These correlated genes were used in a predictive signature to correlate patient outcome with predictive amplification status (Figure S4) revealing that metastasis was associated with the amplification event (Figure 3D).

**Figure 3:**
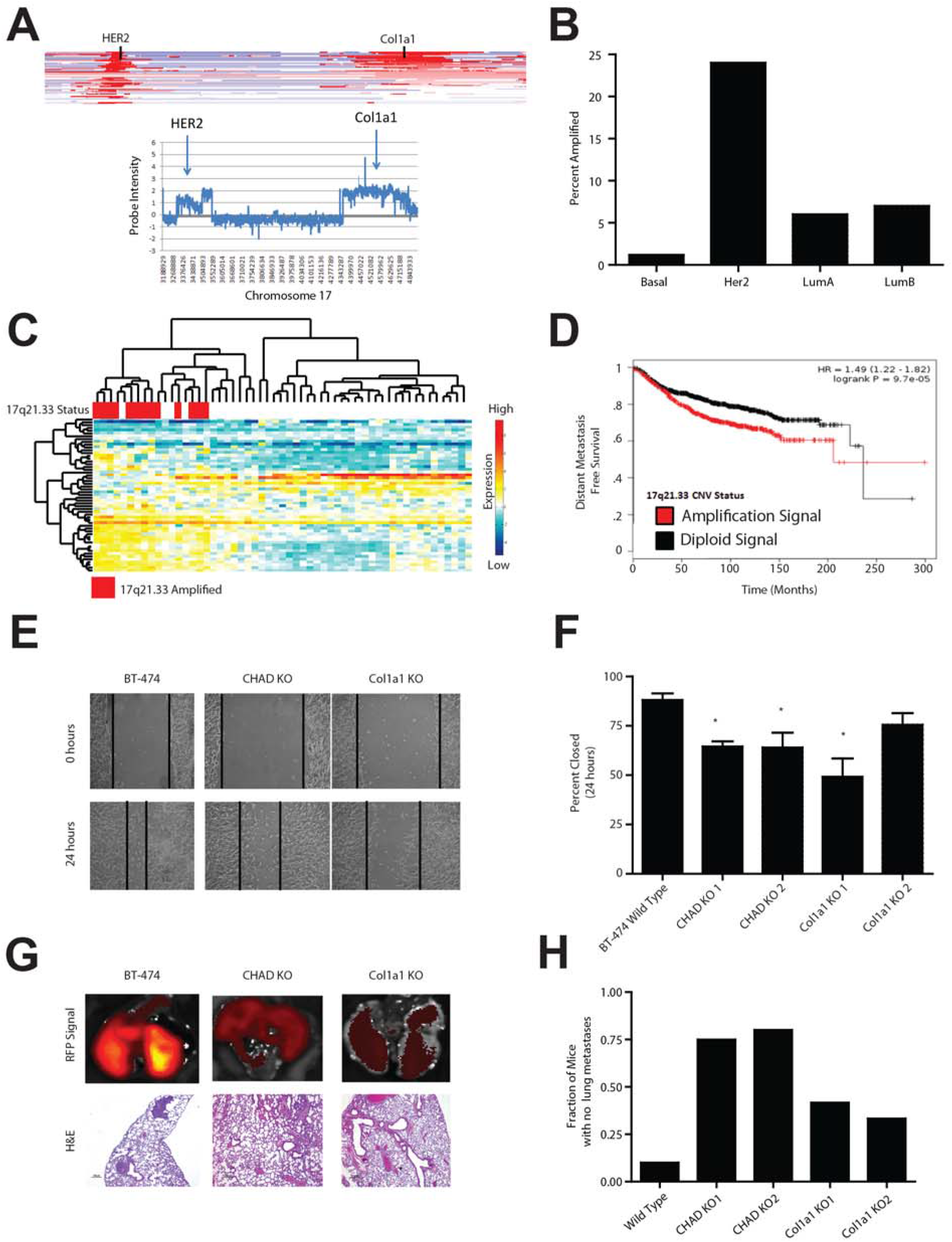
11D amplicon presence and function is conserved in human breast cancer. TCGA breast cancer copy number dataset analysis revealed co-amplification of HER2 and the COL1A1 locus through a heatmap across chromosome 17 of multiple samples with each row representing an independent patient sample (A-top) with red representing amplification and blue representing deletion. The COL1A1 amplification event occurred independently of HER2 (A-bottom) as identified by probe intensity of aCGH data of a single TCGA breast cancer patient. The COL1A1/CHAD amplification event was disproportionately found in HER2 positive tumors and is present in approximately 25% of Her2 positive tumors (B). Gene expression of HER2+ samples with and without the Col1A1 17q21.33 amplification demonstrated a unique gene expression profile as identified by unsupervised hierarchical clustering (C) and overall survival within the KMplotter dataset (P<.001) (D). CRISPRi mediated knockdown of CHAD and COL1A1 in human cell line BT-474 resulted in defects in wound healing (F, G) (*=p<.05, n=9) and distant metastasis to the lung after orthotopic injections (H, I n=10, for WT, n= 4 for CHAD KO1, n = 5 for CHAD KO2 n= 12 for Col1a1 KO1, n=6 for Col1a1 KO2).

To test whether *COL1A1* and *CHAD* were driving the metastasis phenotype in human breast cancer, we used CRISPRi^12^ to knockdown *COL1A1* and *CHAD* in the Her2 amplified, *COL1A1/CHAD* amplified breast cancer line BT-474. These knockdowns showed a decreased ability to migrate in a wound healing assay (Figure 3E and 3F). Importantly the knockdown lines also were unable to metastasize to the lung after being injected into the mammary fat pad (Figure 3G and 3H). Together these data underscore the importance of identifying copy number variation in mouse models of cancer.

In addition to copy number alterations, the whole genome sequence data resulted in the identification of numerous mutations (Figure 1C-E). When COSMIC mutational signatures^13, 14^ were applied to the models, it was observed that the tumor models had similar mutational processes (Figure S5). The MMTV-Neu and MMTV-PyMT tumors both contain the same trinucleotide context of their mutation spectrum. The mutation spectrum shows all nucleotide substitutions present with a slight bias towards C/T and T/C transitions. When compared to the human mutational signatures, the mutational processes present in both mouse models closely resembles COSMIC signature 5 (Fig S5C). This signature has been shown to be present in breast cancer patients with disease associated with late onset^15^, indicating a similar mutational process in both the human disease and mouse models Distribution of SNVs reflected patterns seen in the transcriptional data (Figure 1B) with some events shared between Neu and PyMT tumors while others were unique to the models. Considerable SNV diversity within a model was also prevalent. For instance, the MMTV-Neu model had no genes with shared mutations in all samples and only five genes containing a coding, non-synonymous mutation in more than one sample (Figure S6). Notably we identified mutations within Mucin 4 (*Muc4*) which are potentially impactful due to *Muc4*’s emerging roles in Her2 positive cancer and metastasis^16^. Interestingly, we observed that PyMT induced tumors had more SNVs in the coding regions of the genome. Specifically, these mapped to 34 genes, 9 of which overlapped with Neu tumors. A number of genes with coding mutations specifically in PyMT tumors, including *Matn2*, *Plekhm1*, *Muc6* and *Ptprh* were observed. *Matn2^17^, Plekhm1* and *Muc6^19^* have all been demonstrated to have roles in tumor progression and metastasis and may contribute to the high metastatic capacity of the MMTV-PyMT model.

To test the frequency of these coding mutations in the models as a whole we selected a population of 10 MMTV-Neu tumors and 15 MMTV-PyMT tumors for targeted resequencing. From these tumors we extracted genomic DNA and performed PCR based amplification followed by Sanger sequencing of *Matn2*, *Plekhm1* and *Ptprh*. While *Matn2* and *Plekhm1* confirmed the whole genome sequencing variant calls in the sequenced tumors, additional mutations were not found. Strikingly, *Ptprh* was found to be mutated in 81% of MMTV-PyMT tumors. Furthermore, the *Ptprh* mutation was shown to be homozygously mutated in 21% of PyMT tumors and heterozygously mutated in 60% of PyMT tumors (Figure 4A and 4B). Surprisingly, an identical C to T mutation was observed in each tumor resulting in a valine residue being converted to a methionine at amino acid 483 (V483M). To test for the conservation of mutations of *Ptprh* in mouse strains beyond FVB/NJ, we sequenced *Ptprh* of MMTV-PyMT models in a C57/Bl6, C57/Bl10, CAST, and MOLF backgrounds as well as a different inbred MMTV-PyMT FVB /NJ line. This analysis showed consistent mutation in the structural fibronectin domains (FN3) and the phosphatase domain of *Ptprh* (Figure 4C). Interestingly we found that the two FVB models contained different mutational patterns indicating an impact of environmental and potential epigenetic causes of mutational hotspots.

**Figure 4:**
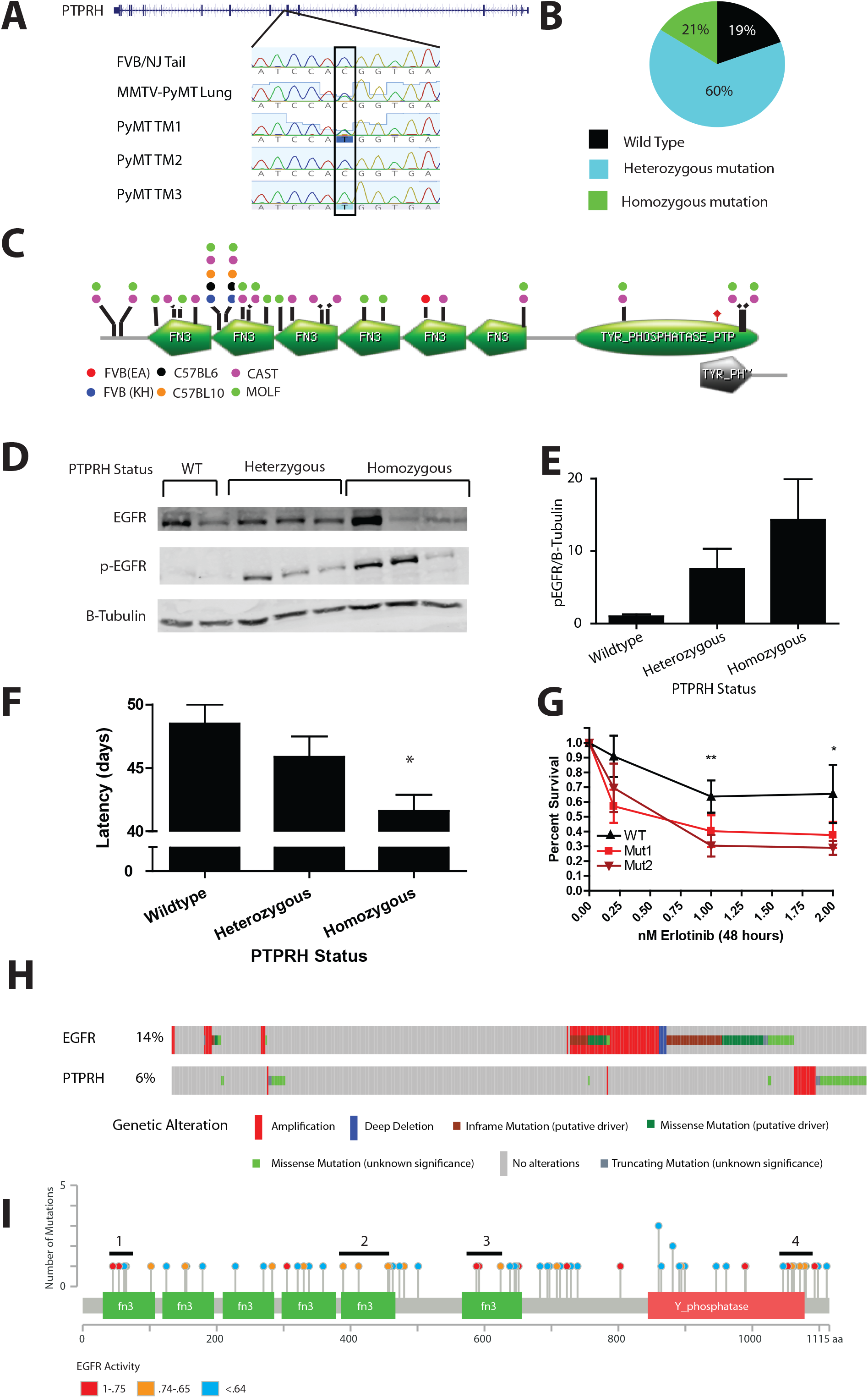
PTPRH mutations are conserved in MMTV-PyMT and human lung cancer. Phosphotyrosine receptor, *Ptprh* was shown through Sanger sequencing (A) to be heterozygously mutated in 60% and homozygously mutated in 21% (B) of MMTV-PyMT tumors (n=45). Sequencing revealed multiple mouse backgrounds have a variety of mutations clustered in the functional domains of the *Ptprh* proteins (C) Increases in pEGFR signaling (C,D) and a decrease in tumor latency (E n=9, *=p<.05) was correlated with mutant *Ptprh* alleles (V483M) within the FVB background. Cell lines derived from *Ptprh* mutant (V483M) PyMT tumors showed an increased response to EGFR targeted therapy including erlotinib (F n=3, **=p<.01, *=p<.05). In human lung cancer (TCGA pan-lung cancer) approximately 5% of patients have a mutation in PTPRH which are mutually exclusive from EGFR (H). High EGFR activity, as determined by gene set enrichment analysis, is associated with mutations clustered within the structural and functional domains of PTPRH as seen by the colors in the lollipop plot (I)

Given that recent work identified the target of *PTPRH* as EGFR^20^, we hypothesized that EGFR was not dephosphorylated with *Ptprh* mutation. Testing this, we observed that the V483M mutation correlated with pEGFR levels (Figure 4D and Figure 4E). With the resulting increase in EGFR activity, we also observed a significant decrease in tumor latency (Figure 4F). With an increase in EGFR activity, it was possible that tumors with mutant *Ptprh* would be dependent upon EGFR signaling. To test this prediction, cell lines derived from *Ptprh* wildtype and mutant tumors were treated with EGFR targeted therapy. After 48 hours, tumors containing *Ptprh* mutations were shown to be more sensitive to erlotinib treatment (Figure 4G).

Given the role of EGFR in lung cancer, we next sought to determine if there was a non-EGFR mutant patient population within lung cancer that could benefit from EGFR inhibition. Examination of the pan–lung TCGA data revealed 5% of patients with a mutation in *PTPRH*. Importantly, these mutations were shown to be mutually exclusive from EGFR, indicating that patients were likely not treated with EGFR tyrosine kinase inhibitors. To confirm the impact of *PTPRH* mutations on EGFR activity in human lung tumors we used gene set enrichment analysis to predict EGFR activity of each mutant *PTPRH* sample. This analysis revealed four key hotspots of mutations driving high EGFR activity, including three in the FN3 domains and one in the phosphatase domain of *PTPRH* (Figure 4I)

Together these data emphasize the heterogeneity within tumor models and the importance of understanding the genomic landscape within the tumors. Here we presented a proof of concept study to identify a number of events that have influenced key tumor phenotypes, including metastasis and tumor latency. Despite the limitations with the number of samples, this study offers a unique opportunity to identify novel genomic alterations which impact tumor behavior and treatment response. These findings have direct therapeutic impact with a potential impact on patient therapeutic intervention and metastatic progression of their disease and underscore the important role genetically engineered mouse models have in understanding tumor biology.

The presence of the *Col1A1/CHAD* amplification event in the mouse model mirrors the 25% of human HER2+ve breast cancers that also had amplification of a structurally conserved region. Given the potential role for these genes in metastasis, this work clearly indicates that the MMTV-Neu system is an appropriate model for select facets of HER2 tumor biology, including the additional amplification event. However, the lack of amplification of genes surrounding *erbB2* in the mouse model indicates that other models with *erbB2* amplification^21^ may be more suitable for other studies.

In the PyMT model system, tumor onset is rapid with mice developing tumors 45 days after birth. Despite this rapid onset, the data presented here indicates that over 80% of PyMT tumors acquire the identical mutation in *Ptprh*, suggesting that there is a significant evolutionary pressure applied during the initial transformation. Given that the *Ptprh* mutant does not dephosphorylate EGFR, this results in unchecked activation of a key signaling pathway. In part, this event may impact metastatic progression of this model system. However, likely the highest impact from this discovery will be in the identification of additional tumor patients that may well benefit from EGFR TKI therapy.

Taken together, this manuscript provides a resource for investigators to determine how well the subtype of cancer they examine is represented by these model systems. While we have explored two genomic events, others of interest are noted and their impact will be elucidated. The data from these two models also underscores the importance of a complete characterization of GEMMs for human cancer.

## Methods

### Animal Studies

All animal husbandry and use was conducted according to local, national and institutional guidelines. The MMTV-Neu^4^ and MMTV-PyMT^5^ mice were in the FVB background. MMTV-PyMT634 and MMTV-Neu mice were obtained from The Jackson Laboratory. Mice were monitored twice weekly for tumor initiation and growth. At a 2000 mm endpoint, mice were necropsied. For mice with multiple tumors the endpoint was established when the primary tumor was at 2000 mm^3^. Tumors and lungs were collected for genomic analysis, hematoxylin and eosin staining for histological subtyping and presence of pulmonary metastases. The number of metastasis was quantified using a single cut through the lung and count of the number of micro-metastases in that plane. Masson’s trichrome staining was used to examine tumors for collagen deposition using standard methods.

### Whole genome sequencing

Flash frozen tumor pieces were ground and DNA was extracted with the Qiagen Genomic-tip 20/G with the manufacturer’s protocol. DNA was sequenced to a depth of 40X with paired end 150 base pair reads on an Illumina HiSeq 2500 using the Illumina TruSeq Nano DNA library preparation.

### Transcriptomic profiling

1 22 23

Transcriptome data for this study was previously published^1, 22, 23^. Data was downloaded from GSE42533 (MMTV-Neu) and GSE104397 (MMTV-PyMT) as.cel files. Affymetrix expression console was used to normalize each individual dataset using RMA normalization. To remove batch effects between datasets BRFM normalization^24^ was performed with standard parameters.

### Clustering

Unsupervised hierarchical clustering was performed using Cluster 3.0 and heatmaps were created using the MATLAB imagesc function.

### Variant calling

Generated.fastq files were assessed for quality control using FASTQC analysis http://www.bioinformatics.babraham.ac.uk/projects/fastqc. Reads were trimmed for quality using Trimmomatic^25^. After trimming, data was reassessed for quality using FASTQC. Then reads were aligned to the mm10 mouse reference genome using BWA-mem^26^. After alignment, base recalibration and pcr induced biases were removed using PICARD tools (http://broadinstitute.github.io/picard). For variant calling we utilized four software packages, GATK^27^, Mutect2^28^, Strelka^29^, and SomaticSniper^30^. To be a legitimate variant we filtered to only those variants called by 3 of the 4 packages. To control for differences in the FVB strain and the mm10 reference genome we used previously published normal FVB tissue (ERR046395)^31^. To call copy number and structural variants we used Delly^32^. For copy number we used default quality control settings and only analyzed those copy number events which had precise boundaries and were larger than 100KB. For translocations we used default quality control setting and precise breakpoints.

### Variant verification and extended tumor panel sequencing

For verification of SNVs we used PCR based amplification followed by Sanger sequencing. For validation of CNVs we used qPCR on the genomic DNA with the Quantabio PerfeCTa SYBR green kit under the manufacturer’s specifications. Primers for PCR and sequencing are listed in Table S7

### Circos visualization

Representative MMTV-Neu and MMTV-PyMT samples were chosen to be displayed as CIRCOS^33^ plots. CIRCOS plots were generated using CIRCOS v 0.69 and SNVs, CNVs, and translocations were mapped according to their location on the mm10 genome.

### Mutation signatures

Due to the low mutational burden of MMTV-Neu and MMTV-PyMT tumors, mutations were combined into a signal analysis for each model. These samples were processed with MutSpec-NMF^34^ for trinucleotide context and comparison to the known human mutation signatures.

### Cell lines

The PyMT 419 cell lines were a gracious gift from Dr. Stuart Sell and Dr. Ian Guess^9^. The NDL2-5 cells lines were obtained as a gift from Dr. Peter Siegel^8^. The BT-474 cell line was obtained from Dr. Kathy Gallo and validated using fingerprinting analysis performed at Michigan State University.

### CRISPR generated knockouts of PyMT 419 and NDL2-5

CRISPR/Cas9 constructs were created to knockout *Col1a1* and *Chad* in PyMT 419 and NDL2-5. Guides were designed and inserted into Px458, obtained from addgene (Addgene #48138) as a gift from Feng Zhang, as previously described^35^. Cells were sorted using FACS technology into single cells and grown into clonal population, then screened for the presence of INDELs using Sanger sequencing. Knockouts were further confirmed for the NDL2-5 lines using western blot. Guide Sequences are listed in Table S7.

### CRISPRi generated knockdowns in BT-474

Knockdowns of Col1a1 and CHAD were created in the BT-474 line using CRISPRi technology. gRNA were cloned into a plasmid containing the gRNA under the control of the U6 promoter (Addgene plasmid #60955)^36^. Lenti virus was created for stable expression of this plasmid and the stable expression of KRAB-Cas9 fusion protein (Addgene plasmid #60954)^36^. Cells were infected with KRAB-Cas9 expression virus first and selected for uptake by puromycin treatment. The stable KRAB-Cas9, BT474 line was then infected with the virus for stable selection of the gRNA for CHAD or COL1A1. These were then sorted using flow cytometery for RFP expression into a pooled population and validated knockdown through western blot. The plasmids used in the part of the project were obtained through Addgene as a gift from Jonathan Weissman.

### Wound healing assay

Wound healing assays were performed similarly for all cell lines in the manuscript. Cells were grown to 100% confluence in a six well plate then a wound was created in the middle of the plate. Cells were allowed to close the wound for 24 hours in the presence of Mitomycin C growth inhibitor then the cells were imaged. Images were quantified for the amount of migration into the wound using ImageJ.

### Tail vein injection

NDL2-5 *Chad* and *Col1a1* knockout cell lines were injected into the tail vein of syngeneic FVB/NJ mice. Cell were suspended in PBS in a single cell population and injected in a single bolus of 500×10^5^ cells in 50uL. Mice were monitored for 9 weeks then euthanized. At this point, lungs were collected and stained with Hematoxylin and Eosin to identify the presence of pulmonary metastases.

### Mammary fat pad injection

NDL2-5 WT and cell lines were suspended in PBS and injected into mammary gland number four in syngeneic FVB/NJ mice as a single bolus of 1×10^6^ cells. The mice were monitored twice weekly until tumors reached an endpoint of 2000 mm in diameter.

BT474 wildype and CHAD/COL1A1 knockout lines were suspended in a 1:1 concentration of matrigel:PBS mixture and injecting into the mammary gland number four in a single bolus of 1×10^6^ cells. Balb/C nude mice were used for these studies. Tumors were monitored until a size of 1000 mm^3^ in diameter. Tumors were then resected and mice were monitored for an additional four weeks. At necropsy lungs were imaged for RFP using the IVIS imaging system and then processed for hematoxylin and eosin staining.

### Human dataset usage

All human datasets used in this study are publically available and noted as used in the manuscript. For genomic alteration frequency the TCGA Breast cancer^10^ and the TCGA-pan Lung cancer^37^ datasets were used. For the expression based survival data the KMPlot.com dataset^38^ was used.

### Western blotting

Western blots in this manuscript were completed under manufacturer’s specifications. Blocking was performed for 1 hour by incubation at room temperature with the LiCor blocking reagents. Western blots were imaged using the LiCor system. The following antibodies were used: COL1A1 (Origene TA309096), CHAD (Abcam ab104757), EGFR (CST D38B1), pEGFR (Invitrogen PA5-37553), HSP90 (CST 4874S), Beta-tubulin (CST 2128S), anti-rabbit secondary (Licor 926-32211), anti-mouse secondary (Licor 926-68070)

### Erlotinib sensitivity assay

Cell lines derived from *Ptprh* mutant and wildtype tumors were seeded at a concentration of 250 cells/mL and subjected to erlotinib treatment for 48 hours with the concentrations stated in the manuscript. Eroltinib was purchased from Cayman Chemical. After treatment with erlotinib or DMSO control, cells were given fresh media to grow for 7 days. Cells were then fixed and stained with crystal violet for counting.

### Data Availability

The datasets generated during and/or analysed during the current study are available in the GEO and SRA repositories

## Acknowledgements

We thank the members of the Andrechek laboratory for helpful discussions. We thank the Michigan State Investigative HistoPathology Laboratory for the assistance with staining. This work was supported in part by Michigan State University through computational resources provided by the Institute for Cyber-Enabled Research.

## Author contributions

JR and EA collaborated on the study conception, design, and interpretation of results. MSU provided annotation for translocations and CAN analysis. YZ assisted with copy number validation. CL, EB, MJ, and WH provided assistance with in vitro experiments. CR, KS, and KH collected samples and performed WES. KH assisted in the writing of the manuscript and WES study design. JR performed all other experiments and drafted the manuscript. All authors have critically read, edited, and approved the final version of the manuscript.

## Supplemental legends

**Figure S1 - Copy number alterations TCGA Breast Cancer Oncoprint**

Oncoprint of the human TCGA Breast cancer cohort (Nature 2012) displaying the alteration of genes altered at a high rate in mouse models with regards to copy number

**Figure S2 - Confirmation of Col1a1 and CHAD knockout in PyMT 419, NDL2-5, and BT-474 cell lines**

Sanger sequencing of CHAD (A) and Col1a1 (B) KO clones revealed the production of indels within the coding sequence of each protein within the PyMT 419 line. This is also the case where multiple different indels were shown in the Col1a1/CHAD amplified cell line NDL2-5 (C). The confirmation of knockdown was completed through western blot for CHAD (top) and Col1a1 (Bottom). The CRISPRi system with guides against early exons of CHAD and Col1a1 was used to generate knockdowns of the respective genes in the human HER2 positive, COL1A1/CHAD amplified line BT474. The efficiency of knockdown in the pooled population was assessed through western blot for CHAD (E) and COL1A1 (F).

**Figure S3 - Addback of Col1a1 and CHAD in PyMT 419 cell lines**

The addback of wildype *Col1a1* and *Chad* into the CRISPR generated knockout lines showed partial recovery of movement in a scratch assay (*=P<.05).

**Figure S4 - Validation of COL1A1/CHAD amplicon gene expression signature**

A score between 0 (diploid) and 1 (amplified) was generated for the predicted presence of the COL1A1/CHAD amplification event based upon a weighted gene expression data. This signature showed a robust prediction of the amplification event in both the training HER2 positive dataset (A) and the Luminal A validation cohort (B)

**Figure S5 - Mutational Signatures of MMTV-Neu and MMTV-PyMT Models**

The trinucleotide context of MMTV-Neu (A) and MMTV-PyMT (B) samples are similar. They show the presence of every mutation possibility with the overrepresentation of the C>T and T>C transitions. These trinucleotide signatures were compared with human mutational signatures through the use of a Baysian model high similarity (Red) and low similarity (yellow) were identified through the use of a heat map. Signature 5 represented the highest similarity score.

**Figure S6 - Heterogeneity of SNVs in mouse models of breast cancer**

The MMTV-Neu (A) and MMTV-PyMT (B) models have considerable diversity in regards to SNVs. Samples were analyzed for overlap in SNV calls through the use of a Venn Diagram

**Table S1 - Tumor characteristics**

A table listing the characteristics of tumors used in this study including, model, latency, histological subtype, and number of lung metastases

**Table S2 - Validation of bioinformatic calls**

The table contains the PCR and qPCR validation of the bioinformatic calls of SNVs and amplification events. Alterations labeled as WT were not identified as altered in copy number or SNV by the variant callers. These were tested to control for false positives.

**Table S3 - Single nucleotide variants**

A table listing called SNVs within the sequenced MMTV-PyMT and MMTV-Neu samples. This table also contains the predicted impact of the SNVs on the protein content and function

**Table S4 - Copy number alterations**

A table listing called copy number alterations within the sequenced MMTV-PyMT and MMTV-Neu samples

**Table S5 - Translocation events**

A table listing called translocations within the sequenced MMTV-PyMT and MMTV-Neu samples

**Table S6 - COL1A1/CHAD amplicon gene signature**

A list of gene names and weights that correlate with Col1a1/CHAD amplification generated through WGCNA analysis which are used to create the predictive Col1a1/CHAD amplification gene expression signature

